# *C. elegans* exhibits coordinated oscillation in gene activation in single-cell developmental data

**DOI:** 10.1101/114074

**Authors:** Luke A.D. Hutchison, Bonnie Berger, Isaac Kohane

**Affiliations:** Massachusetts Institute of Technology, 77 Massachusetts Ave, Cambridge, MA, USA.; Harvard Medical School, 25 Shattuck St, Boston, MA, USA.

**Keywords:** *C. elegans*, body plan, development, developmental clock, developmental age, transcriptional regulation, single-cell, gene ontology, anatomy ontology, Fisher’s Discriminant

## Abstract

**Background:** The advent of *in vivo* automated single-cell lineaging and sequencing will dramatically increase our understanding of development. New integrative analysis techniques are needed to generate insights from single-cell developmental data.

**Results:** We applied novel meta-analysis techniques to the EPIC single-cell-resolution developmental gene expression dataset for *C. elegans* to show that a simple linear combination of the expression levels of the developmental genes is strongly correlated with the developmental age of the organism, irrespective of the cell division rate of different cell lineages. We uncovered a pattern of collective sinusoidal oscillation in gene activation, in multiple dominant frequencies and in multiple orthogonal axes of gene expression, pointing to the existence of a coordinated, multi-frequency global timing mechanism. We developed a novel method based on Fisher’s Discriminant Analysis (FDA) to identify linear gene expression weightings that are able to produce sinusoidal oscillations of any frequency and phase, adding to the evidence that oscillatory mechanisms likely play an important role in the timing of development. We cross-linked EPIC with gene ontology and anatomy ontology terms, employing FDA methods to identify previously unknown positive and negative genetic contributions to developmental processes and cell phenotypes.

**Conclusions:** This meta-analysis demonstrates new evidence for direct linear and/or sinusoidal mechanisms regulating the timing of development. We uncovered a number of previously unknown positive and negative correlations between developmental genes and developmental processes or cell phenotypes. The presented novel analysis techniques are broadly applicable within developmental biology.

## Background

Four-dimensional microscopy (imaging three dimensions over time) and a range of single-cell sequencing and profiling techniques are enabling unprecedented study of how the developmental program unfolds *in vivo* [1, 2, 3, 4, 5, 6, 7, 8, 9]. However, as the amount of available single-cell data increases, we need better tools for analyzing and visualizing these datasets (e.g. [10]), in order to make the data more comprehensible and useful.

One of the early, groundbreaking whole-organism 4D confocal microscopy techniques, by Bao, Murray, Waterston *et al.*[5, 7, 8, 11] employed histone-mCherry reporters under the control of upstream promoters to detect expression levels of genes of interest at single-cell resolution, while employing fluorescent cell labeling to enable automated cell lineaging. This process yielded a complete three-dimensional map of gene expression at single-cell resolution across the entire developmental timeline while simultaneously tracing the cell division pedigree, resulting in a 4D gene expression dataset superimposed over the cell division tree. This dataset was published as EPIC, Expression Patterns In Caenorhabditis [12]. Subsequent research has applied a number of other techniques to the problem of building single-cell profiles during development (e.g. [9]), but the EPIC dataset remains uniquely interesting and valuable, since the dataset tracks expression levels of over 125 developmentally-interesting genes at single-cell resolution from the zygote throughout early development, while also automatically tracing the cell pedigree.

Despite evidence that a global biological clock may govern the fate of cells and the timing of development, mechanisms regulating the timing of development that have been discovered so far appear to be localized, and a comprehensive control mechanism for global developmental timing has yet to be determined [13, 14, 15, 16, 17, 18, 19, 20, 21, 22]. Recent work on single-cell gene expression has illuminated interesting regulatory processes at work during development [23, 24, 25, 26, 27, 28], but until recently, there has been comparatively little data available on the expression patterns of expression across multiple genes, in all cells of an organism, across the entire developmental timeline. As a range of experimental techniques have recently become available for producing single-cell profile data across the entire development timeline of an organism [9, 29, 30, 31], new methods are needed for determining the mechanisms behind the regulation and timing of development, utilizing the data produced by single-cell techniques.

Despite evidence that a global biological clock may govern the fate of cells and the timing of development, mechanisms regulating the timing of development that have been discovered so far appear to be localized, and a comprehensive control mechanism for global developmental timing has yet to be determined [13, 14, 15, 16, 17, 18, 19, 20, 21, 22]. Recent work on single-cell gene expression has illuminated interesting regulatory processes at work during development [23, 24, 25, 26, 27, 28], but until recently, there has been comparatively little data available on the expression patterns of expression across multiple genes, in all cells of an organism, across the entire developmental timeline. As a range of new techniques have recently become available for producing single-cell profile data across the entire development timeline of an organism [9, 29, 30, 31], new methods are needed for determining the mechanisms behind the regulation and timing of development, utilizing the data produced by single-cell techniques.

Here, we present a meta-analysis of a dense subset of the EPIC dataset that newly provides evidence that a comprehensive control mechanism governs development in *C. elegans*. This subset consists of the expression levels of 102 developmentally-relevant genes across the first 686 cell identities in the *C. elegans* cell pedigree (i.e. all ancestral cell identities up to approximately the 350-cell stage). We did not include genes with low coverage over these core cell identities, or cells beyond this point for which there was little gene expression data. Significant work was required to prepare the EPIC data for meta-analysis (Methods). Producing the complete gene expression matrix for these genes and cells allowed us to apply novel analysis techniques to the data.

We applied principal component analysis to the EPIC gene expression data for *C. elegans* and newly observed gene expression following a sweeping manifold shape through expression eigen-space, as the developmental timeline proceeded (Figure 1). We discovered a strong linear correlation (*R*^2^ = 0.94) between the first principal component of gene expression and wall-clock time during cell proliferation (Figure 2). Over the entire available timeline, we observed multiple apparent sinusoidal oscillations in gene expression (Figure 3), with different frequencies of oscillation manifest in different principal components, indicating that our observation of linear monotonic correlation between gene expression and wallclock time have been an observation of the nearly linear part of a sinusoidal graph around the zero-crossing point (it is difficult to know without the availability of data from later in development). Whether what we observed is linear or sinusoidal in nature, what we observed in the first principal component is remarkable, in the sense that a simple linear projection of gene activation patterns moves monotonically in a specific direction, in some low-dimensional basis embedded in the high-dimensional space of gene expression. This is strong evidence for a the global coordination of development as a choreographed, chronological sequence – in other words, it is evidence that a global clock regulates the entire developmental process.

**Figure 1.**
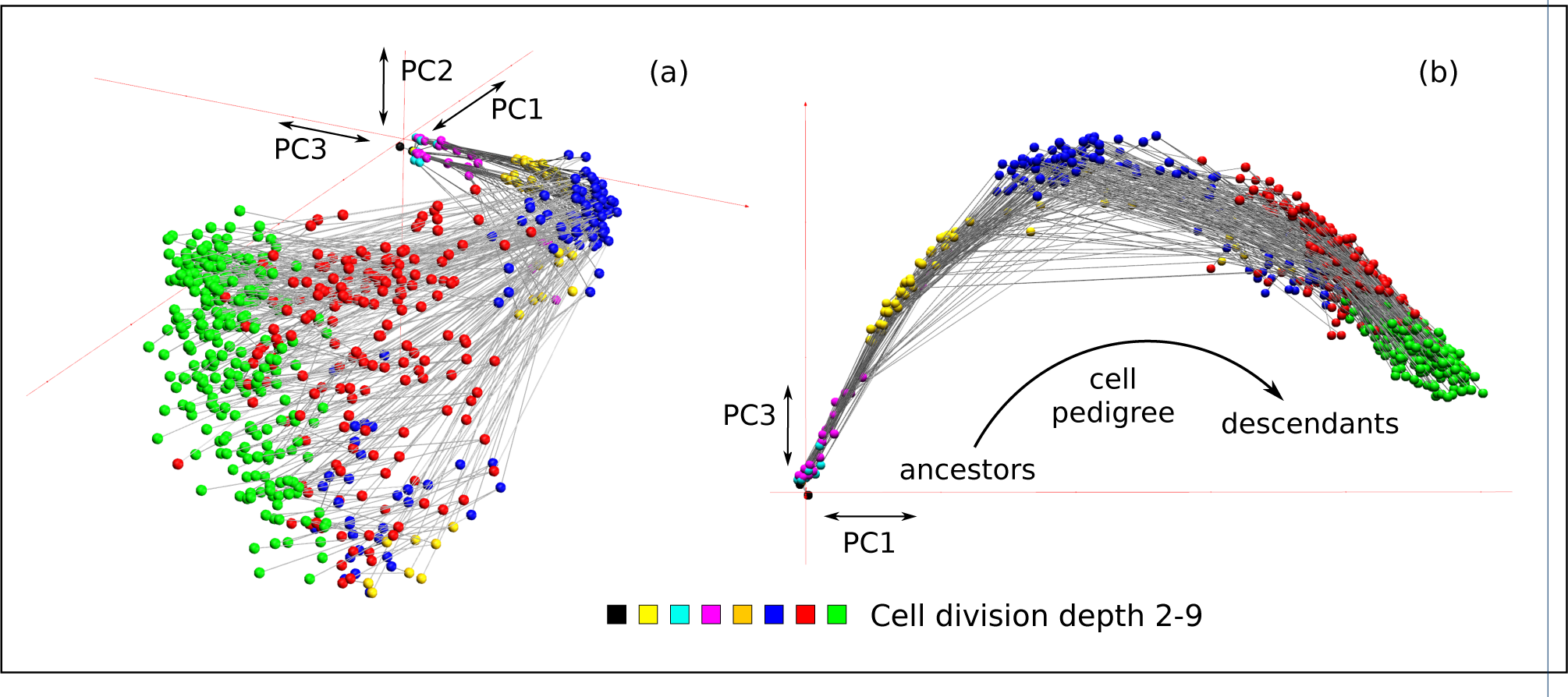
Projection of binarized gene expression profiles onto the first three principal component axes. Each node represents a cell, and the edges between the nodes connect a cell with each of its two daughter cells. The color of each node indicates the division depth in the cell pedigree. (a) A perspective view of the projection of the data onto the first three principal components PC1-PC3, showing that the cell pedigree is embedded in a curved manifold that sweeps through space in a direction primarily aligned with PC1 as development proceeds. (b) A top-down view of PC3 vs. PC1, showing the curved path of the manifold relative to the first principal component axis.

**Figure 2.**
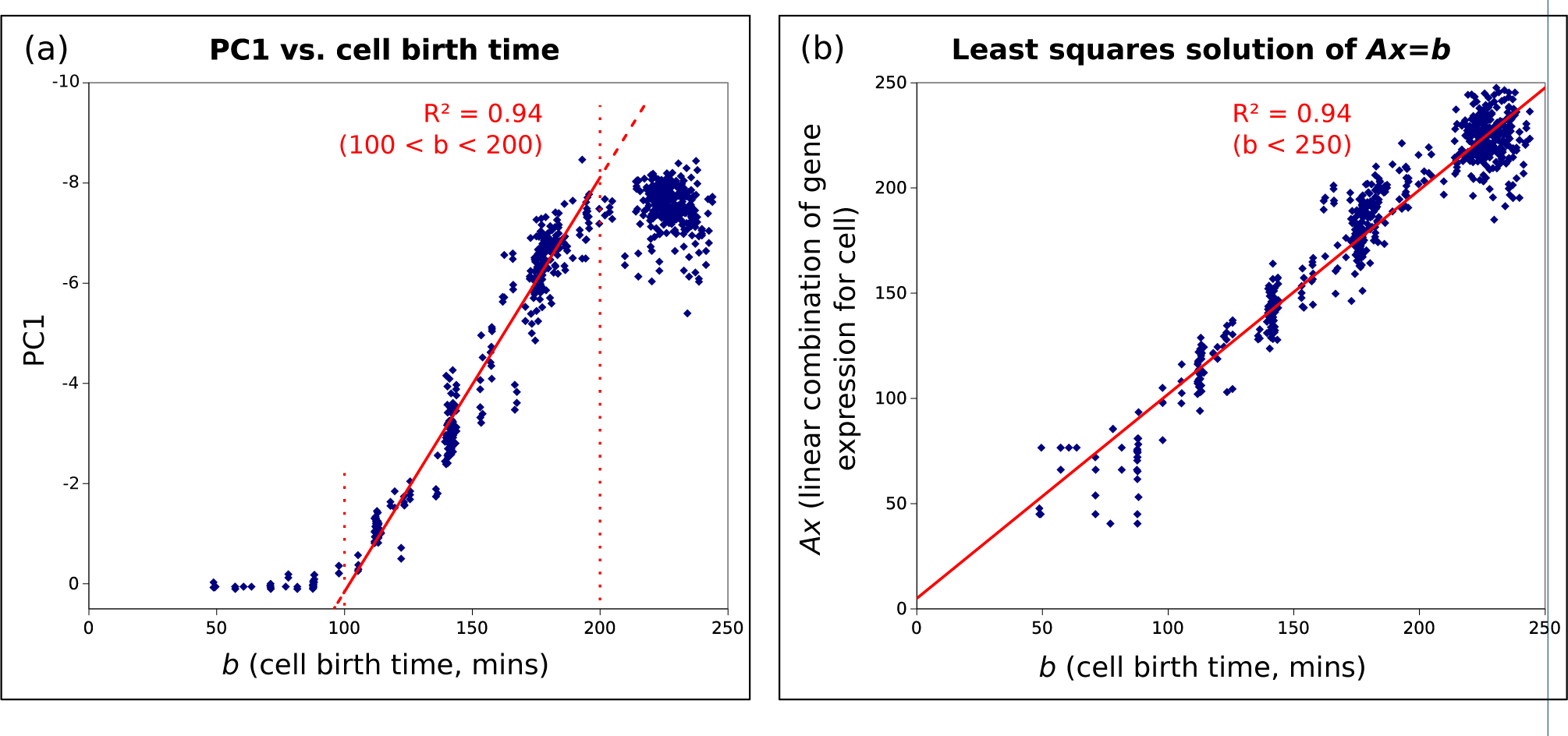
(a) the projection of gene expression onto the first principal component, PC1, vs. *b*, the cell birth time (i.e. the cell onset time) in minutes. The plot is strongly linear from 100 to 200 minutes. (b) After solving the linear equation ***Ax = b***, where ***A*** is the binarized gene expression matrix and ***b*** is the vector of cell birth times, this plot shows the components of ***Ax*** (the best-fit linear estimators of each component of ***b***) against the corresponding component of ***b***. The model is linearized across the entire developmental timeline, using a fixed linear combination of gene weights.

**Figure 3.**
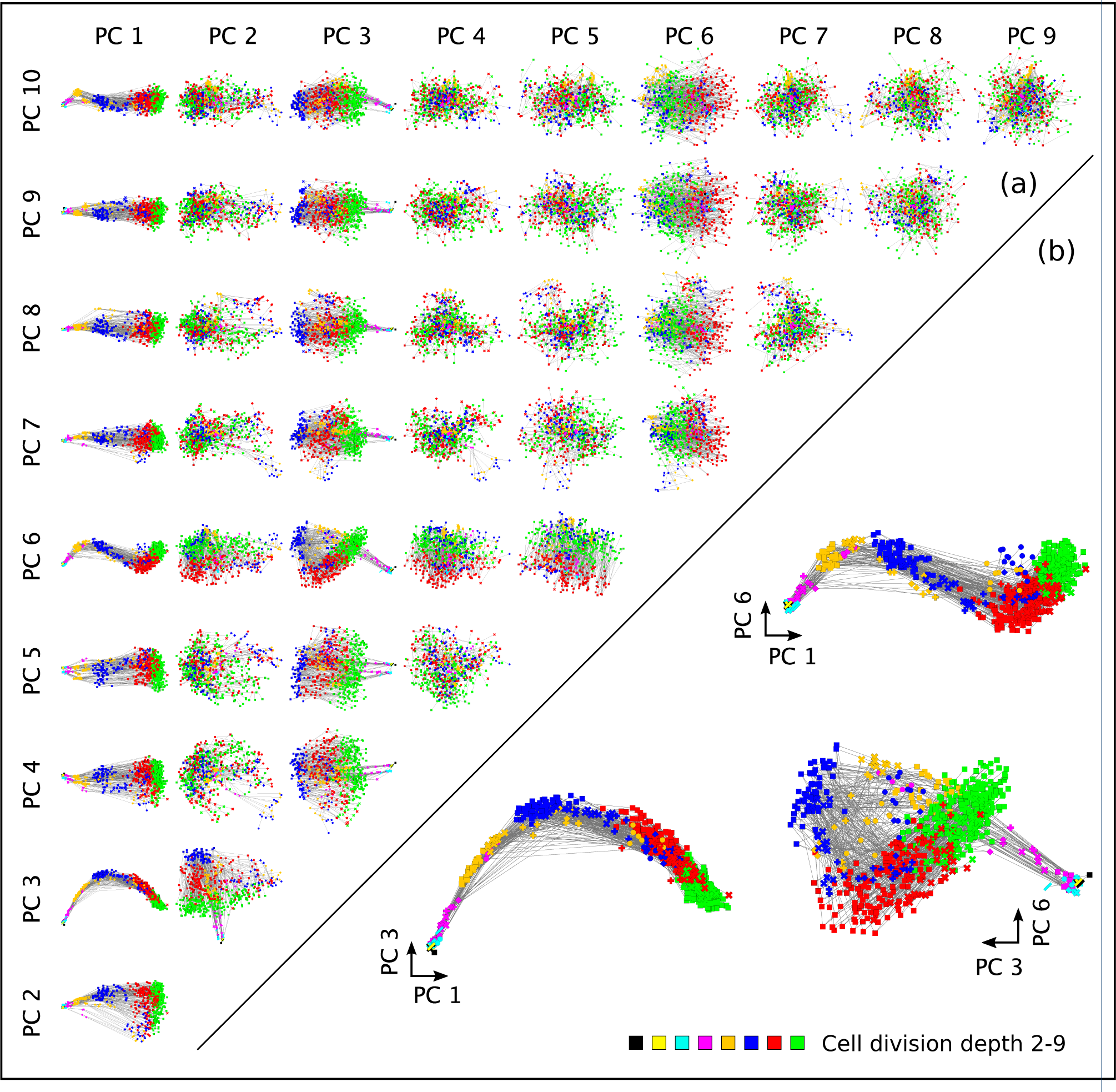
(a) The projection of gene expression onto the first 10 principal component axes, PC1-PC10, shown as 2-dimensional projections: the row and column labels indicate the pairing of two component axes that produces a given projection. (b) Expanded views of PC3 vs. PC1, PC6 vs. PC1, and PC6 vs. PC3. Gene expression appears to oscillate sinusoidally in PC3 and PC6, with approximately double the sinusoidal frequency in PC6 (although insufficient data was available to determine if either PC3 or PC6 would continue a sinusoidal path later in development). Plotting PC6 against PC3 causes the cell pedigree to trace an “alpha”-shaped path (α) through gene expression space as development proceeds.

We devised a novel technique from Fisher’s Discriminant Analysis (FDA) for uncovering the relative contributions of genes to an attribute of interest. It is a powerful yet simple mechanism to determine which genes most strongly contribute (positively or negatively) towards a phenotypic trait or developmental process of interest, by splitting cells into two groups, one with the trait and one without, then finding the set of gene weights that maximizes inter-class variance while simultaneously minimizing intra-class variance (Figure 4).

**Figure 4.**
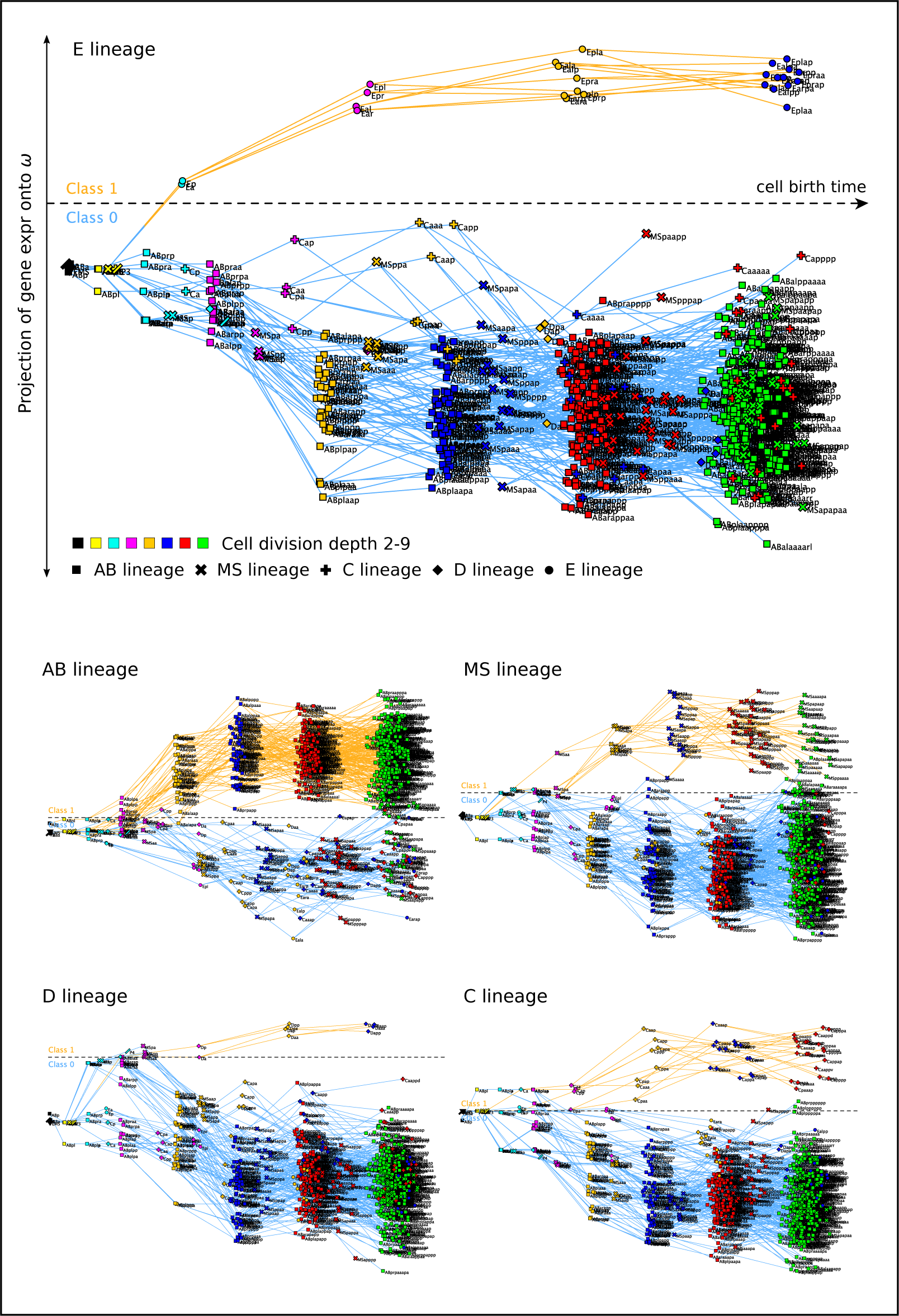
Separation of the E cell lineage (Class 1) from other cells in the cell pedigree (Class 0) using Fisher’s Discriminant Analysis (FDA). The horizontal axis indicates developmental time, and the vertical axis represents the linear projection onto the one-dimensional vector ω that gives maximal inter-class variance and minimal intra-class variance. The resulting gene weights are strongly positive for genes primarily expressed in the E lineage, and strongly negative for genes primarily expressed in cells other than those in the E lineage.

To test the extent to which our observation of multiple clear sinusoidal modes of oscillation may pervade the dataset in other frequencies and phases not observed in the basis axes of the top few principal components, we employed the FDA technique we developed to find linear weightings of gene activation that were able to reproduce sinusoidal oscillations of any desired phase or frequency (Figure 5). This result suggests that indeed, oscillatory mechanisms may be used extensively to regulate the timing of development, through, in the most simple case, a positive or negative linear dependency upon the expression levels of genes of interest.

**Figure 5.**
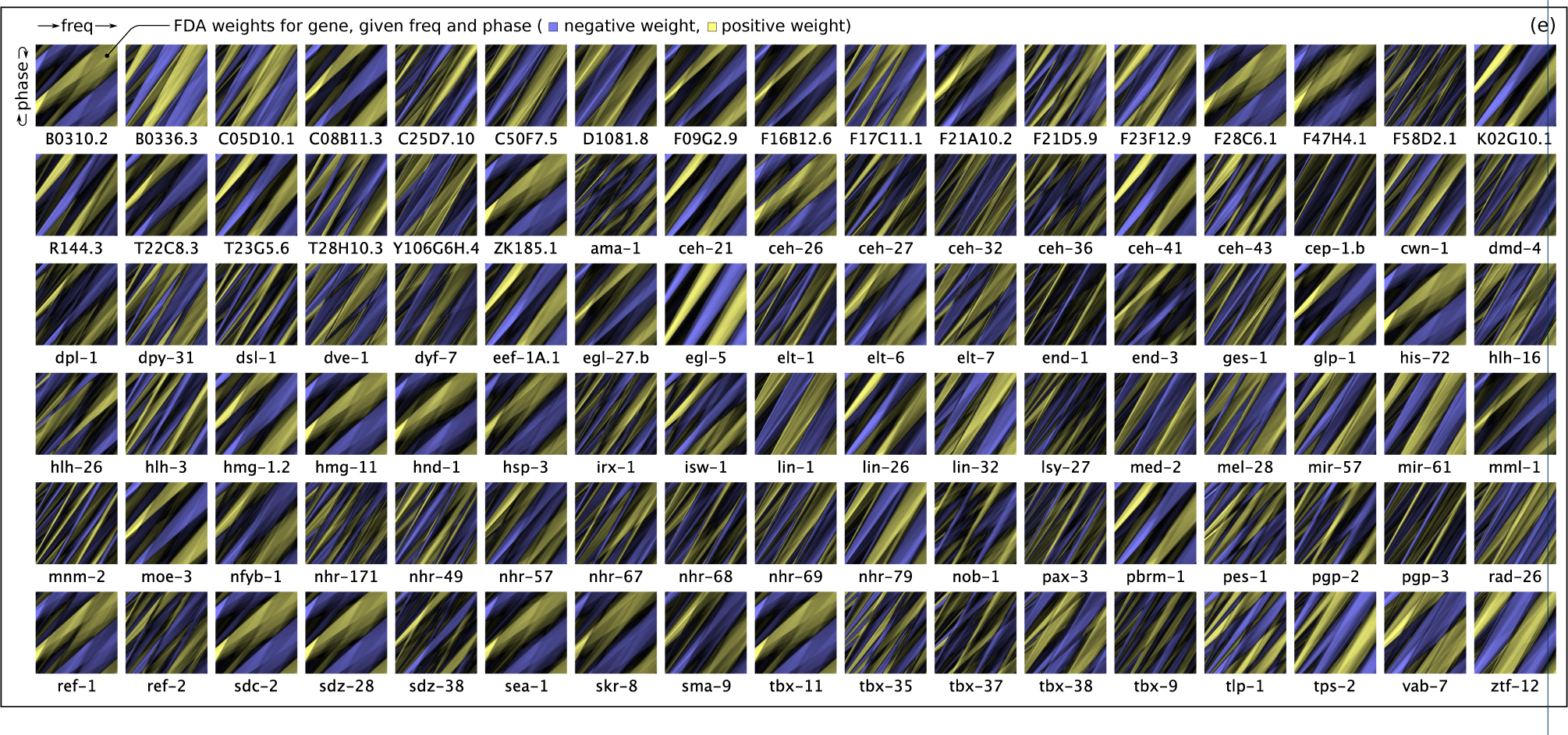
Identification of simple linear weightings of gene expression levels that can produce oscillations across a range of sinusoidal frequencies and phases. (a) A target sine wave is generated. (b) Cells are assigned to class 0, at developmental times when the sine wave is positive, or class 1, at times when the sine wave is negative. (c) Fisher’s Discriminant Analysis (FDA) is used to maximally separate class 0 from class 1 in the vertical axis, minimizing intra-class variance and maximizing inter-class variance, producing a best-fit square wave approximation of the target sine wave. (d) The best-fit FDA results are plotted across a range of phases in the rows and frequencies in the columns, with phase wrapping vertically (0 → π → 0), and with frequency increasing across the columns. (e) A heatmap of FDA weight given phase and frequency for each gene, with the largest negative weight for the gene in blue, and the largest positive weight for the gene in yellow.

The presence of whole-organism sinusoidal and/or linear trends in gene expression would suggest the existence of a global, coordinated mechanism regulating developmental timing, in the form of a monotonic clock, and/or one or more global oscillatory mechanisms. This global timing mechanism may either involve the genes directly observed in this dataset, or may be due to an unobserved, external mechanism acting upon these genes.

To determine the functionality of developmental genes of interest, we also applied our FDA technique to the cross-linked EPIC dataset with the gene ontology and anatomy ontology from Wormbase to identify weightings for each gene that specify how strongly a gene appears to be correlated with the presence or absence of a given phenotypic trait or developmental process in each cell. These gene weightings provided a large number of previously unknown implications about the functioning of various genes across a wide range of developmental processes.

The methods described in this paper are simple to apply, but have the potential to be broadly useful in understanding single-cell experimental data. The specific results we produced from our meta-analysis of the EPIC *C. elegans* data, cross-linked with Wormbase ontologies, presents strong evidence for global control of developmental timing, suggesting numerous opportunities for further research into the time-correlated and oscillatory mechanisms of developmental regulation.

## Results

### Most variance in gene expression is characterized by the first 3-10 principal components

We produced a two dimensional matrix of gene expression in the EPIC dataset, with the 686 *C. elegans* cell identities (up to the 350-cell stage of development) in the rows, and the 102 developmentally-relevant genes in the columns. Each gene was marked as either active in the cell (1.0), for fluoresence levels above 2500, or inactive (0.0), for fluorescence values below this level. To examine the contribution of the principal axes towards overall dataset variance, we produced a *scree plot* (Figure S1) from the binarized gene expression matrix (Table S1), showing the eigenvalues of gene expression sorted into decreasing order. Most of the variance in the expression patterns of the 102 genes is embodied in the first 10 principal components, and in particular by the first three principal components.

### The cell pedigree monotonically sweeps a curved manifold through gene expression space

To understand how patterns in gene expression changed in relation to cell division, we produced a novel three-dimensional visualization of the cell pedigree directly overlaid on the first three principal components of gene expression (Figure 1). In this plot, nodes represent the 686 unique cells in the dataset (specifically, the specific identities of cells between cell division events), and edges indicate the relationship between a cell and its two daughter cells. The color of a cell in this plot indicates the cell division depth. The position of a cell in the three-dimensional space is the projection of that cell’s gene expression profile (the binarized expression levels of the 102 genes for the cell) onto the first three principal components, i.e. the cell’s position in this 3D plot is a simple linear combination of the activity levels of the cell’s genes. By definition, the first principal component (PC1) is aligned with the axis of greatest variance in gene expression, the second principal component (PC2) is aligned with the axis of second greatest variance in gene expression orthogonal to PC1, and the third principal component (PC3) is aligned with the axis of third greatest variance in gene expression orthogonal to both PC1 and PC2.

Because the edges that connect each non-leaf cell to its two daughter cells clearly show the cell pedigree, trends in gene expression can be clearly seen as development proceeds. Remarkably, the cell pedigree sweeps across a curved manifold surface embedded in the three-dimensional “eigengene” expression space. The sweep direction of the cell pedigree across the manifold is monotonic, in the sense that pedigree edges between cells and their daughter cells all follow the same approximate sweep direction; there are no pedigree edges directed opposite to this general sweep direction after the first two or three cell divisions.

The strongest vector component of this sweeping manifold path is aligned with the first principal component of gene expression (PC1), indicating that movement along the manifold in the direction of the pedigree is monotonically correlated with the most significant orthogonal direction of variance in gene expression. The implication of this is that the largest variation in gene expression across all cells is time-correlated, and this time correlation is monotonic – in other words, variations in gene activation is collectively coordinated and *sequenced* such that development moves in a specific direction.

The strong, monotonically smooth manifold shape collectively swept out by the expression patterns of these genes suggests a strong and tightly coordinated global gene expression control mechanism. Since a large number of genes are collectively involved in sweeping this broad path through gene expression space (as seen in the principal component weightings, Figure S4), this could imply that the developmental program is redundantly encoded across many genes for robustness, or alternatively, that many of the genes studied are directly or indirectly regulated by a global timing mechanism, or both.

As cells divide during development, and as gene expression trends in the direction of PC1, the cloud of cells at a given cell division depth (indicated by a given node color) expands in width along PC2 relative to the previous cell generation. This behavior indicates a general diversification in gene expression patterns as development proceeds, consistent with lineage-specific differentiation. However, the total spread in PC2 is less than half the distance swept through PC1 as development proceeds, suggesting that variation in expression levels of genes in this dataset is more strongly associated with the progress of development than the details of tissue-specific differentiation, since PC1 is most strongly aligned with the developmental timeline.

### Gene expression is correlated with developmental age of the organism

Remarkably, since each cell pedigree edge from a cell to its two daughter cells follows the arrow of time, and since the cell pedigree sweeps a “monotonic” manifold shape through gene expression space as development proceeds, the expression patterns of the selected genes must be related to the developmental age of the organism, or at least to the cell division depth. Figure 2(a) shows PC1, the first principal component of gene expression, plotted against *b*, the birth time of each cell in minutes since fertilization. For much of the recorded development time, a striking linear correlation is evident (*R*^2^ = 0.94 from 100-200 minutes). This correlation is notable, because projection of the data onto the PC1 axis represents a simple weighted linear combination of the binarized gene expression levels, which indicates there is a weighting of the gene expression levels that is directly predictive of the wall clock developmental age in minutes. This linear weighting can be obtained directly from the eigenvector loadings for the first principal component (i.e. the eigenvectors multiplied by the square root of the corresponding eigenvalue – Figure S4; Table S2).

This strong correlation between PC1 and developmental age appears to be independent of the cell division rate, in particular implying that the gene expression profiles of cells are all similarly correlated with developmental age regardless of cell division depth within the cell pedigree. This finding can be seen by the skew in cell division depth in Figure 1, evidenced by the increased mixing of cell colors, representing cell division depth, as development proceeds. Each of the major cell lineages MS, C, D and E have successively slower cell division rates relative to the AB lineage (the difference in cell division rates between the lineages can be seen in Figure 4, but gene expression is not correlated with cell division depth as strongly as with developmental age). The finding that wide-scale gene expression patterns are decoupled from cell division rate agrees with the findings in Nair *et al.* [22], who found that expression and proliferation are independently entrained to separate clock-like processes. Whereas Nair *et al.* found that the *relative timing* of cell division was not directly correlated with large-scale transcriptional regulation, in our analyses we observed that the *depth* of cell division was not directly correlated with large-scale transcriptional regulation, since lineages with dramatically different cell division intervals exhibit similar expression trends.

Before 100 minutes and after 200 minutes, however, PC1 diverges from being linearly correlated with the cell birth age *b* (Figure 2(a)). It is possible gene expression is linearizable across the entire developmental timeline, but that PCA does not recover the optimal set of gene weights to expose the direct linear correlation, and for this purpose we consider below a linear regression method for fitting a linear model to the data.

Another possible explanation for the observed distribution is that the underlying gene expression is fundamentally sinusoidal, not linear, and that we merely observed the section of the sinusoid that is most linear, around the zero crossing point (discussed below). It is also possible that entirely different developmental programs or processes are active before 100 minutes, between 100 and 200 minutes, and after 200 minutes: one plausible explanation for the difference in gene activity before and after 100 minutes is the maternal-to-zygotic transition (MZT) [32].

To find the optimal function mapping gene expression to cell birth time, we could try using supervised learning with randomized k-fold cross validation to learn a regression function. However, this would tend to overfit the data and underestimate the prediction error, because so many cells share similar gene expression profiles (meaning that there would be sample pollution between the training set and the test set). Instead, we expressed the relationship between gene expression and cell birth time as a linear system ***Ax*** = ***b***, where ***A*** is the binarized gene expression matrix (i.e. Table S1, with cells in the rows, and genes in the columns) and ***b*** is the vector of birth times for each cell. We then solved the system for ***x***, a set of gene weights that map the expression matrix onto cell birth times. (A small amount of random noise was added to the binarized gene expression levels for regularization.) The resulting vector of gene expression weights ***x*** (Table S5) gives us the linear weighting of genes that maps the gene expression profile of a cell onto the developmental age of the cell with the minimum squared error.

Given this set of weights, gene activity was close to linearly correlated with cell birth time for all cells in the pedigree (not just from 100 to 200 minutes), retaining approximately the same linear correlation strength of *R*^2^ = 0.94, but across the entire developmental timeline (Figure 2(b)). If PC1 is evidence of a linear correlation with developmental age, rather than a sinusoidal correlation, then the genes with high-magnitude positive or negative weights in this table would be indicated as important in the timing of development. The gene with the largest negative weight in Table S5, *med-2,* is necessary for endoderm specification [33]. Other genes with high negative weights include *sdz-28*, an SKN1-dependent zygotic transcript active only in early development, triggered by the SKN1 maternally-deposited transcription factor, and *glp-1*, which encodes a transmembrane protein essential for mitotic proliferation of germ cells and maintenance of germline stem cells, and important in many differentiation decisions in somatic tissues. Genes with strong positive weights include *egl-5*, a Hox gene [34], as well as a number of genes that are expressed nearly ubiquitously across the entire developmental timeline.

Table S6 lists, for each cell, the cell birth time, the PC1 projection of gene expression for the cell, and the linearized gene expression for the cell, and compares these with the Sulston onset time of the cell.

### Sinusoidal oscillation observed in the principal components of gene expression

Figure 1 shows that gene expression in the third principal component, PC3, traces half a sinusoidal cycle with respect to PC1. The Waterston technique [5, 7, 8] is not able to reliably track development once the organism begins to move, so the end of the current dataset is not the end of the developmental timeline. It is unclear whether a complete sinusoid would be traced in PC3 if more of the developmental timeline were captured – in other words, it is not clear whether the semi-sinusoidal oscillation in PC3 relative to PC1 is in fact half of a sinusoidal oscillation. Figure 3(a) shows all pairings of principal components between PC1 and PC10 as a mechanism of visualizing the first ten principal component dimensions in 2D, and interestingly, PC6 traces a complete sinusoid with respect to PC1 across the same time period that PC3 traces half a sinusoid. If PC6 is paired with PC3, the half sinusoid paired with the complete sinusoid causes the developmental path to trace a spiraling alpha shape (α), as shown enlarged in Figure 3(b). The presence of a complete and clear sinusoidal oscillation in PC6 relative to PC1 lends strength to the hypothesis that the semi-sinusoidal curve of PC3 relative to PC1 is in fact half of a sinusoidal oscillation.

Principal components other than PC1, PC3 and PC6 did not exhibit strong linear or sinusoidal structure, but may capture variation in gene expression due to tissue-specific variation. The widening of the point cloud in all principal components other than PC3 and PC6 when plotted against PC1 is consistent with cells differentiating as development proceeds. See for example the widening in PC2 vs. PC1 in Figure 1.

The semi-sinusoidal oscillation observed in PC3 and the sinusoidal oscillation observed in PC6 may also explain the nonlinearity observed in plotting PC1 against developmental age (Figure 2(a)) before 100 minutes and after 200 minutes – the plot overall exhibits a sigmoidal shape. This suggests that in fact PC1 may be oscillating sinusoidally over the developmental timeline, and that the observed linearizable portion of PC1 between 100 and 200 minutes is in fact only linear because a sinusoidal wave is also nearly linear around its zero-crossing point. Given more data about later stages of development, sinusoidal oscillation may be exhibited for all three of PC1, PC3 and PC6, with periods of roughly 500 minutes for both PC1 and PC3, and 250 minutes for PC6. If these oscillations are in fact sinusoidal, then given that the phases of the apparent oscillations in PC1 and PC3 are offset by π*/*2 from each other, and that they have similar periods, the plot of PC1 vs. PC3 (Figure 3(b)) should trace a roughly circular motion.

### Fisher’s Discriminant Analysis used to identify lineage-specific genes

It is possible to determine a linear combination of gene expression levels that maximizes the separation between a group of cells that possess a specific trait (Class 1) and cells that do not possess the trait (Class 0), using Fisher’s Discriminant Analysis (FDA), a specific, statistically-grounded method for Linear Discriminant Analysis (LDA). The FDA method finds the one-dimensional vector ω which, when the dataset is projected onto it, maximizes inter-class variance and minimizes intra-class variance (Methods). Using the FDA method, we obtained gene weightings that maximally separate each major lineage in the *C. elegans* pedigree – AB, MS, C, D and E – from cells in other lineages (Figure 4). In this figure, the projection of gene expression onto ω is used as the y-axis value for each cell, and the birth time of each cell is used as the x-axis value. The clear separability of each cells in each lineage from the remainder of the cells indicates that gene expression patterns in each lineage are distinct.

Table 1 lists the gene weights required to produce maximum class separation in the projection of the gene expression of cells onto ω for the E lineage. Genes with weights that are more strongly positive are generally expressed in the E lineage and not expressed in other lineages, whereas genes that are strongly negatively weighted are generally strongly expressed in cells other than those in the E lineage. This finding allows for the identification of strongly lineage-correlated genes. Note however that this is somewhat different than just looking to see which genes are preferentially expressed in a given lineage – this is a weighting of gene expression levels that most cleanly separates the E lineage from other cells, and that produces the most compact separate clustering of the E lineage and non-E lineage cells. This result is also different from what would be obtained with PCA, and in general, PCA axes are not aligned with LDA axes, meaning that they do not maximally separate distinct clusters or classes: PCA identifies axes of greatest overall variance, whereas LDA (or FDA) finds the axis of greatest class separation. Note that *e*nd-1, the top-weighted gene in the FDA weightings for the E lineage, is well known as an important developmental gene for the E lineage (the intestine) [35]. Heat shock-driven expression of END-1 has been found to cause a majority of embryonic cells to express intestine-specific genes and form intestinal structures [36]. It has also been discovered than mutations in the second-highest ranked gene, nob-1, along with pgp-3 (ranked in the top 25% of gene weights for the E lineage), both posterior-group Hox genes, results in gross posterior embryonic defects [37]. However, not all of the highly-weighted genes have been previously linked to intestinal development in *C. elegans*. More importantly, the *negatively weighted* genes for the E lineage or most other developmental processes, body structures, or phenotypic traits, have rarely been examined. This table raises the question as to whether strongly negatively-weighted genes such as isw-1 (Chromatin-remodeling complex ATPase chain) are specifically not expressed in the E lineage – but similar questions could be asked about negative gene weights for each cell lineage, or each cell attribute, tested using FDA.

**Table 1.**
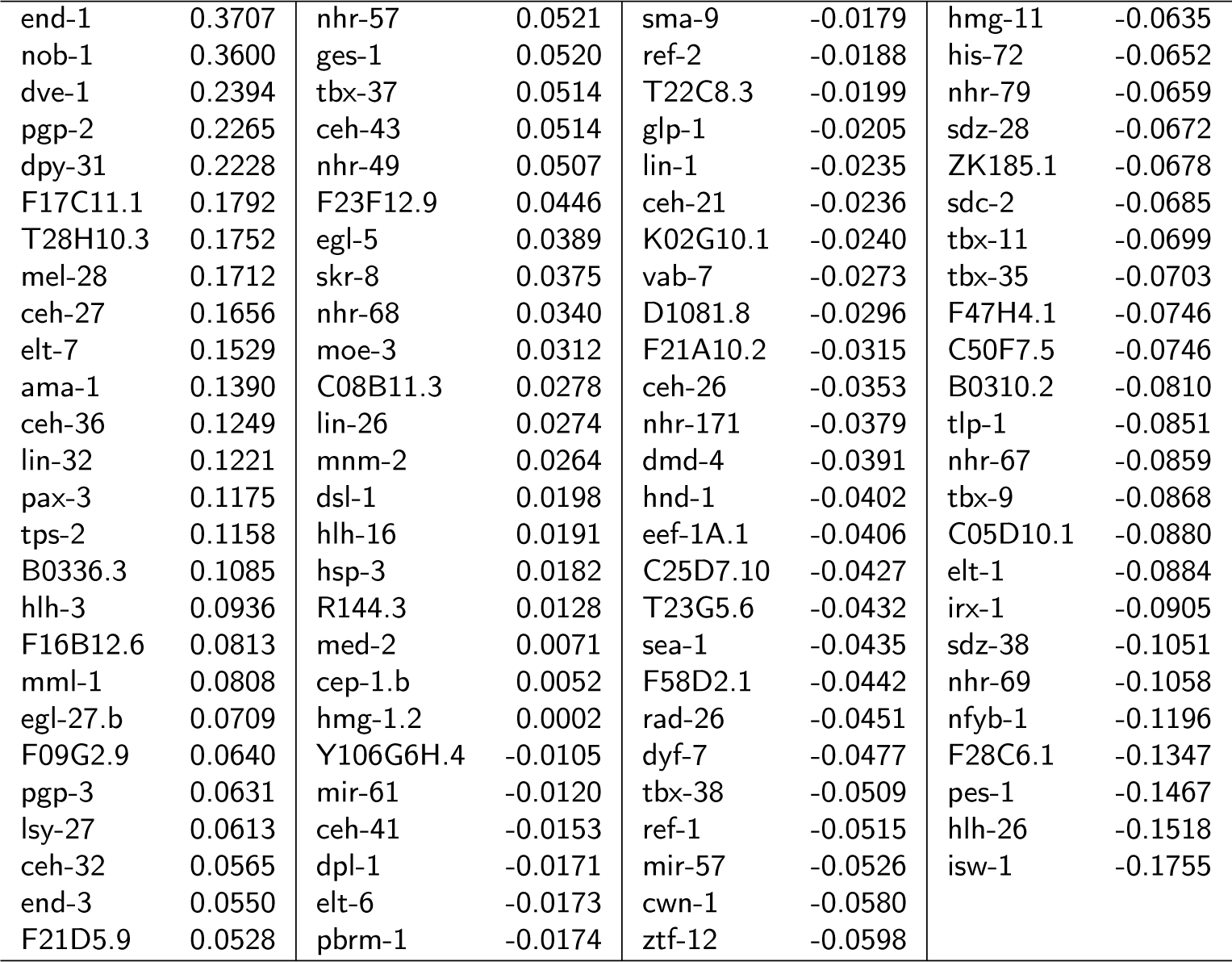
Gene weights for FDA separation of E lineage from other cells. Strongly positive genes are predominantly expressed in the E lineage; strongly negative weights are predominantly expressed in other lineages. The FDA process that produces these weightings maximizes inter-class variance (variance in gene expression between the E lineage and other cells) while minimizing intra-class variance.

Of note, *elt-1*, which has been identified as a master regulator of epidermis specification, is strongly weighted in separating the AB epidermal lineage from the rest of the cells, but not as strongly weighted in separating the C epidermal lineage from the rest of the cells (Table S4). This is consistent with the observation that these two lineages rely on different developmental regulators [38]. Also, several factors are weighted more highly than *elt-1* in separating the AB lineage from the rest of the cells (*sdz-38*, *tlp-1*, *hlh-26*, *hlh-3*, *pax3*), and several factors are weighted similarly highly in separating the C lineage from the rest of the cells (*nob-1*, *cwn-1*, *C25D7.10*, *tbx-9*, *rad-26*). Some of these genes are known to play a central role in epidermal development, including *nob-1* [39], *pax-3* [40], and *tbx-9* [41], confirming the utility of this FDA method for identifying genes relevant to developmental processes. However, the other highly-weighted factors for the AB and C lineages do not appear to have yet been closely studied in relation to their broader role in epidermal development. The same is true of the highly weighted genes for each of the major lineages – some but not all of the highly positively-weighted genes have documented roles in the development of these lineages, and, importantly, most of the strongly negatively-weighted genes have not previously been identified as specifically being inactive only in a given lineage.

### Insights into roles of genes in development

We next sought to identify overall patterns in the weightings of each gene in the FDA results, given known functions of each gene. To this end, we crosslinked the FDA results with the Wormbase Gene Ontology terms (GO terms) for each of the 102 genes under study. Given a set of gene weights, we looked up the GO terms for each gene, and contributed the weight of the corresponding gene into an accumulator for the GO term. The sum of the gene weights for all genes associated with each GO term, for the E lineage FDA weightings, are presented Figure 2. These aggregate weightings give an idea of which biological processes are more characteristically active during intestinal development compared to the development of the rest of the organism. There are RNA polymerase II terms at both the high positive and low negative ends of the scale, which could be related to inconsistent application of GO terms to genes in the GO database, or the inconsistent application of multiple related and similar but differently-coded terms. Either way, there are numerous strong developmental signals at the positive and negative ends of the scale, as would be expected, and specifically, “endodermal cell fate specification” is predictably highly-weighted for gut development (the E lineage). Other terms stand out as interesting, such as “nematode male tip morphogenesis” – indicating that development of the male tail tip is coordinated with development of the gut, or that development is at least regulated by some of the same genes in both cases.

Note that Figure 4 and Tables 1 and 2, constitute the FDA results for just one attribute, where Class 1 is comprised of cells in the E cell lineage and Class 0 is comprised of all other cells in the cell pedigree. For comparison, we also applied FDA analysis to each of the other major lineages in *C. elegans* (AB, MS, C, and D) vs. the other cells in the pedigree (also visible in Figure 4), and found that all the major lineages were cleanly separable from cells not in that lineage, indicating that each lineage had a distinctive gene expression profile. We also tested a couple of hundred other cell attributes, derived from Wormbase anatomy ontology terms, including tissue type and cell function, as well as division depth, and other attributes. The full set of FDA result can be seen in Table S4.

**Table 2.**
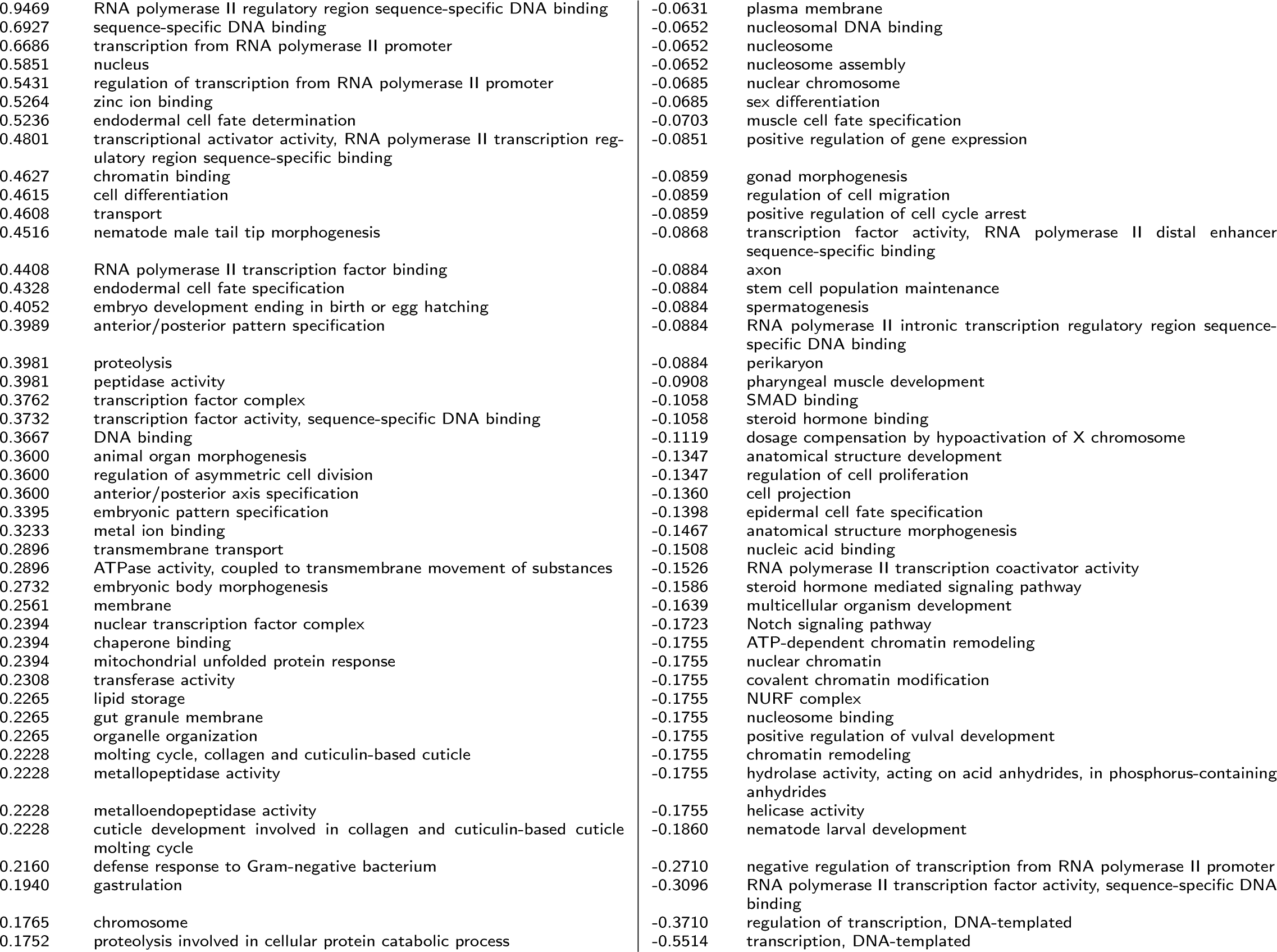
The largest positive and negative gene ontology term (GO term) scores for FDA separation of E lineage from other cells. (GO terms of intermediate magnitude are not shown.) These scores are calculated by summing the position of a cell in its projection onto the FDA ω vector for all cells labeled with a given gene ontology term. Strongly positive gene ontology scores are predominantly associated with cells in the E lineage; strongly negative gene ontology scores are predominantly associated with cells in other lineages. As expected, endodermal cell fate determination scores highly for the E lineage (0.5236).

We also applied the FDA technique to a number of cell features, including anatomy ontology terms from Wormbase, as well as developmental stage, tissue types, etc. For each, we generated a figure indicating the separation of cells that were labeled with the trait from cells that were not labeled with the trait, as well as FDA gene weightings and GO term weightings for each FDA result (Table S4). We also produced variants of each case, for cells that were parent cells or daughter cells of cells that possess each given trait. In each case, we measured the ratio of inter-class to intraclass variance, as a measure of separability of the two classes. This ratio is given in the filename of the FDA results.

### Identification of gene weights producing arbitrary sinusoidal oscillations

The PCA analysis we performed showed that a simple linear combination of gene expression weights (the principal component axes) exhibited apparent sinusoidal oscillations of three different phases and frequencies. In order to determine the extent to which other oscillatory gene activation patterns, may be exposed by simple linear weightings of gene activation, beyond those evident in the first few principal components, we examined the extent to which simple linear combinations of gene expression could produce sinusoidal oscillations of any desired frequency and phase. In other words, given a desired target frequency and phase of oscillation, was it possible to find a linear combination of gene activations that would produce that oscillation? Finding a generic mechanism for producing gene expression oscillations of any frequency and phase could be evidence that sinusoidal oscillations of a wide range of frequencies play a ubiquitous role in the timing of developmental events, and that any mechanism dependent upon an oscillation of specific frequency and phase could obtain that oscillation as a simple linear function of the activation patterns of a subset of genes.

We temporally divided cells into two classes based on the cell birth time in the developmental timeline, corresponding to peaks (Class 0) or troughs (Class1) in the desired sinusoid (Figure 5(a)). Fisher’s Discriminant Analysis was used to find the hyperplane that separates class 0 and class 1 with minimum intra-class variance and maximum inter-class variance, which effectively created a square wave approximation of the target sine wave. The projection of gene expression onto the normal vector ω of the separating hyperplane was plotted against the cell birth time to produce a best fit of gene expression to the original sinusoidal wave (Figure 5(b, c)).

Across a wide range of different phase and frequency values, this method was able to produce a reasonable approximation of a sinusoid (Figure 5(d)). This is a strong indicator that oscillatory behavior of almost any required frequency or phase could be obtained as a simple linear combination of the expression levels of these genes. Figure 5(e) shows the FDA weights for each gene that are required to achieve maximum class separation at a given frequency and phase. The largest positive and negative FDA weights in each gene’s heatmap correspond to the gene’s largest contributions to separating class 0 and 1 at a given frequency and phase (Discussion).

### Analysis of transpose PCA

For reference, Table S7 gives the PCA of the transpose of the gene expression matrix, which shows the result of dimension reduction in the cell dimensions rather than the gene dimensions. The distance between any pair of genes in this transpose-PCA space, calculated using the Euclidean norm on the top *k* principal components (e.g. for *k* = 2), gives a measure of the similarity of expression patterns for the pair of genes. For example, *sdc-2*, *hmg-11*, *F28C6.1*, *tbx-11*, *B0310.2* and *ceh-26* are close together in transpose-PCA space (their first two transpose PCA coordinates are very similar), and the gene expression patterns for those genes are similar, as shown in Figure S3, whereas *egl-5* is distant from all other genes in transpose-PCA space, and has the most unique gene expression profile, also shown in Figure S3.

## Discussion

### Connection to previous observations of oscillatory patterns during development

Circular oscillatory paths in gene expression have been previously observed with PCA dimension reduction on whole-organism RNA-Seq profiles in frog, mosquito, fly, and zebrafish, by applying the Traveling Salesman algorithm to a series of RNA-Seq profiles to arrange the samples into approximate order of developmental age, finding a minimum-distance simple path between all samples [42]. This is significant, as it lends credence to the presence of a global clock mechanism coordinating development. The expression levels of individual mRNAs were also observed to oscillate over time [43, 32]. Both oscillatory and temporally gradated activity has been observed in transcript levels [44]. Some key regulators of the timing of heterochronic miRNA expression have been discovered, including *lin-4* and *let-7*-family miRNAs, under control of *lin-42* [45], however data on these miRNAs was not available in the EPIC dataset for comparison.

Robust oscillation in transcription has been observed previously at multiple temporal scales [43, 44], involving ultraradian cycles affecting approximately 1*/*6th of the transcriptome, with changes in expression of up to an order of magnitude during the cycles. The authors identified several periodic developmental phenomena, such as cuticle development and cuticle molting, and speculated that the timing of other developmental processes are similarly controlled by one or more of these transcriptional cycles.

What is unique about our findings is that the patterns of oscillation in gene expression that we observed were not limited to specific cell types, or to specific cell lineages, and were not affected by the cell division rate of different lineages. Our observations were not made by taking samples of transcripts or other genetic activity, averaged across the whole organism at specific timepoints. Rather, we observed oscillations occurring *separately and simultaneously within each individual cell at single-cell resolution*, as part of a globally synchronous oscillatory pattern exhibited by all cells in existence at each point in the developmental timeline of the organism. Also notable is our observation of *multiple superposed oscillations of different frequency and phase*. This superposition of time-correlated oscillation was *contemporaneous with non-oscillatory patterns of gene expression involved with cell differentiation*: the observed patterns of gene activation simultaneously and collectively encoded multiple oscillatory mechanisms, in an almost “holographic” sense, based on gene activations at individual cells – and yet each gene simultaneously and separately also served its own unique role in development, unrelated *per se* to the oscillation to which it contributed.

### Possible conflation of similarity in cell function with co-temporality

In a wide array of gene expression research, in order to gain insights into the functioning of genes, cells have been clustered according to similarity in their gene expression profile (e.g. [4]). However, Figure 1 illustrates a possible important caveat to understanding these clustering results: cells with the most similar gene expression profiles may be more strongly *temporally* related than they are *functionally* related. This is true at least of the dataset we studied: the greatest source of variance in gene expression that we observed (PC1 in our PCA results) was *time-correlated*. This implies that, at least for the genes selected for this dataset, *differential gene expression is more strongly affected by developmental age than it is by tissue-specific differences in expression arising from differentiation processes*.

Note however, as a caveat, that in correspondence, Murray [7] pointed out that the genes selected for analysis in the EPIC dataset may have been biased for temporal patterns, as opposed to spatial patterns. Therefore, a larger study, involving a wider array of genes, would be needed to determine whether or not the observed effect was due to the selection of genes.

If however the above observations are generally true of larger sets of genes, across longer spans of the developmental timeline, then many previous research conclusions about cell clustering and cell similarity may need to be revisited in the light of how gene expression is globally coordinated across all cells at each stage of development, for example by the oscillatory processes we observed in PC3, PC6, and PC10, and in particular, by the highest-variance component, the monotonic progress of the gene expression manifold through gene expression space that we observed in PC1. The magnitude of the correlated variance observed in these principal components cannot be discounted in the analysis of cell similarity based on gene expression similarity alone.

### Cylindrical projection of gene expression manifold

The manifold swept by the cell pedigree through the space spanned by the first three principal component axes was roughly semi-cylindrical. We flattened out this “principal manifold” of the data [46] using a cylindrical projection. This involved radially projecting cell positions in principal component space outwards from a central axis onto the surface of a cylinder, and then flattening out the cylinder (Figure S2). Table S6 gives the two dimensional coordinates (θ, y) of the cells in the cylindrical projection.

This cylindrical projection can be used to visualize the cell pedigree in two dimensions rather than three, while eliminating most problems with occlusion and perspective distortion. We plotted the binarized expression data for each gene using the cylindrical projection in Figure S3, so that large-scale patterns of gene expression can be examined across the developmental timeline, and across the surface of the principal manifold traced through the first three principal components.

### Examination of principal component weights

The principal component weights (i.e. the gene weights that project the gene expression data onto the principal component axes) indicate which genes are the strongest sources of variance in a given principal component axis (Table S2(b); Figure S4). If variance in a principal component axis is only due to a small number of genes, the weights corresponding to those genes will be large in positive or negative magnitude, and the other genes will be close to zero. However, the distribution of PC1 weights in Figure S4 demonstrates that the contribution towards variance is close to zero for very few of the 102 genes in this dataset, indicating that most or all of the 102 genes under consideration are involved in establishing the linear correlation with developmental age. If the time-correlated nature of PC1 is indeed due to a global developmental clock mechanism, then the fact that many genes are involved in this mechanism could indicate a redundancy in the temporal functioning of these genes, affording the opportunity for adaptation in the function of developmental regulators without disrupting the global developmental clock. Redundancy adds resilience and flexibility, giving a system more degrees of freedom over which to adapt.

However, comparing the sorted weights in Figure S4 to the gene expression patterns in Figure S3, it can be seen that the strongest negative weight (*egl-5*) corresponds to gene expression in all cells excluding the last generation measured, whereas the strongest positive weights (*lin-26*, *K02G10.1*, *ceh-41*, etc.) correspond to high levels of gene expression commencing later during development. Consequently, the sorted PC1 weights roughly correspond to a time ordering of gene activation. It is possible then that the apparent time-correlatedness of PC1 is due to a sequential pattern of gene activation. However, cross-comparison with Figure S3 suggests that the situation is not as simple as the genes being switched on in a specific ordered sequence.

### Generation of sinusoidal oscillation as a linear weighting of gene activations

In Figure 5(d), we showed that a simple linear combination of gene activations could produce a sinusoidal wave of any desired frequency or phase. This result lends support to the idea that the patterns of sinusoidal oscillation that we found in our principal component analysis, as a simple linear projection onto the principal component axises, were not an anomaly, but may have been due to an underlying process that relies upon the combined activation of positively weighted genes and the combined inactivation of negatively weighted genes to measure oscillation during development, with several important frequencies being tracked by the developmental processes of the organism. The fact that we could recreate this phenomenon for any desired frequency or phase may indicate a deeper pattern – that oscillatory timing of developmental processes as a simple function of gene activations may be a mechanism that is relied upon ubiquitously during development.

One possible explanation for the ability to generate oscillations of any phase and frequency lies in the fact that the different genes studied become active and inactive at different times during development (Figure S3). As long as the predominant timespans of the activity of different genes do not perfectly overlap, an appropriate weighting of genes could select a subset of genes that demonstrate activation during the peaks of a sine wave, and do not demonstrate activation during the troughs of a sine wave, as a function of time. This raises a “chicken or egg” question about whether sinusoidal oscillations may control gene expression, or whether gene expression collectively gives rise to sinusoidal oscillation. Perhaps both processes are happening, and are interrelated. This question deserves further research.

### Effects of gene selection and reporter mechanism on results

Gene activity data was collected by Murray *et al.* using the following criteria and methodology: “We identified a list of transcription factors and other regulatory proteins for which prior microarray or phenotype data suggested embryonic function and targeted these for expression analysis. For these, we constructed stable *C. elegans* strains expressing a histone-mCherry reporter under the control of the gene’s upstream intergenic sequences. We analyzed expression of reporter strains whose expression begins before the last round of embryonic cleavage (the 350-cell stage) by crossing in a ubiquitous histone-GFP marker, collecting threedimensional confocal time-lapse movies, and tracing the cell lineage as described previously.” [7]

In correspondence, Murray suggested the use of histone-mCherry reporters may impact analysis, because these reporters tend to persist and even increase in the descendants of expressing cells, even if the endogenous gene is degraded (the reporter mRNA has a “stable” *let-858* 3’ UTR, and the histone-mCherry itself has a halflife that is likely to be longer than the length of embryogenesis). It is unclear what the effect of gene selection and reporter mechanism may be on our results, and further work is needed to determine whether other gene sets and/or different reporter methods for obtaining single-cell-resolution gene expression data exhibit the same properties.

## Conclusions

We have presented a comprehensive meta-analysis of the *C. elegans* single cell resolution EPIC gene expression dataset of Waterston *et al.*. Our analyses show multiple temporal patterns in the expression data, including oscillatory and/or linear correlations vs. developmental age, hinting at a global regulatory mechanism or developmental clock. We show that a simple linear weighting of gene expression can be chosen to exhibit roughly sinusoidal oscillation of any desired phase or frequency, suggesting that sinusoidal oscillations may be pervasive in regulating development. These results warrant further study, in order to understand and characterize the mechanisms of global regulation of gene expression during the developmental process. We presented a number of novel techniques, and novel applications of existing statistical techniques to whole-organism developmental data, which yield novel insight into regulatory genes and regulatory control mechanisms during development. These techniques should be broadly applicable to similar single-cell or whole-organism datasets.

## Methods

### Dataset preprocessing

We obtained the Waterston EPIC dataset from [12]. This dataset comprises image data (obtained using 4D confocal microscopy of developing *C. elegans* embryos) as well as gene expression data for 127 developmentally related genes at single-cell resolution up to the point at which the embryo gains motor control.

We parsed the available data, discarding genes and cell pedigree subtrees with significant numbers of missing values (i.e. where gene expression had not been recorded for significant numbers of cells). For genes that were run multiple times, we averaged the expression levels across the runs.

We took the maximum expression level of each gene across the lifetime of each cell, using the “global” intensity measurement method (out of “global”, “local”, “blot” and “cross”, as described in the original paper), and binarized gene expression to 0 or 1 using a reporter intensity threshold of 2500. This threshold value was conservatively chosen based on Murray *et al.* [7], where it was stated that spurious gene activity was not observed below a measured reporter intensity value of 2000, and that strong expression signals were observed at values over 4500. We discarded a number of additional genes that were expressed in all (or nearly all) cells, as well as genes that were not expressed in any (or almost any) cells given this intensity threshold.

The resulting binarized gene expression matrix (Table S1) consists of the binarized gene expression values (0 or 1) for 102 genes, measured in 686 cells. The 686 cells can be broken down into 341 internal nodes in the cell pedigree (cells that divide within the measured developmental timeline) and 345 leaf cells (cells that are either terminally-differentiated, die through apoptosis, or divide later than the end of the recorded timeline).

### Timescale correction

We extracted cell birth times from the dataset by finding the time point for each cell at which reporter intensity level data first became available (these times were used as the x-axis for the linear regression in Figure 2). Despite the claim in [7] that the EPIC data was sampled “with ∼1-min temporal resolution”, the raw data was actually sampled at different time scales for each gene, with time scale factors including at least 1.0, 1.35, 1.4, 1.5 and 2.0 minutes per 3D scan, and with the datasets listing only scan indices, not timestamps. There is no available data source on the EPIC website that indicates the time scale for a given run, and in private correspondence, the original authors were not able to easily retrieve the timescales used for each run. Therefore, to get all data on the same timescale, we had to perform some slightly tricky analysis to recover a best-fit time scale factor for each run. We used a custom multiple-alignment regression technique to warp the gene expression timepoints for each run to a consensus cell pedigree. In this process, we also discovered that not only a multiplicative offset was needed to scale the data indices to fit the timeline of the canonical Sulston pedigree, but there was also an additive offset averaging approximately 45 minutes between the Sulston time zero and the dataset index zero (i.e. the sample indices in the raw datafiles are zero-indexed, but the sampling started at a developmental age of approximately 45 minutes). Cell birth times, appropriately scaled and offset as described above, were used for the linear correlations in Figure 2.

Note that despite our best efforts to align these datasets, a small degree of time-spreading has probably been unavoidably been introduced into our cell birth time predictions, due to the fact that the raw data was not properly timestamped. Properly timestamping future scans using minutes since time of fertilization (rather than sample index) would increase the strength of linear correlation (*R*^2^) between developmental age and gene expression.

The EPIC data includes multiple runs for some genes, sometimes with nontrivial variation in gene expression between runs. After adjusting for the unspecified time dilation as described above, we combined data from these different experiment repetitions by averaging, across all runs for a gene, the maximum reporter level achieved by the gene during the lifetime of each cell.

### Principal component analysis

Principal component analysis was run on the binarized gene expression matrix, producing Table S2(a) and Table S2(b), the PCA eigenvalues and eigenvectors respectively. We projected the binarized gene expression matrix onto the eigenvectors, producing Table S3.

The figures in this paper were produced using our own custom 2D and 3D visualization software. Columns 1-3 of Table S3 were plotted in a 3D view to yield Figure 1. All pairings of principal component axes for PC1-PC10 were plotted as 2D projections, yielding Figure 3.

### Cylindrical projection

The cylindrical projection of gene expression in Figures S1 and S2 were produced by identifying a central axis in the 3D-embedded two-dimensional manifold in which lies the cell pedigree, as shown in Figure 1. This central axis was chosen such that cells in Figure 1 were all approximately equidistant from this axis. The rotation about this axis (θ) and position along the axis of the line segment orthogonal to the axis that passes through each cell (y) were used as 2D coordinates for the cylindrical projection plot.

### Fisher’s Discriminant Analysis for identifying genetic basis of differentiation

Figure S4 and Figure 5 were generated by implementing Fisher’s Discriminant Analysis (FDA). This method employs Fisher’s linear discriminant to provide a closed-form solution for maximizing inter-class variance while minimizing intra-class variance between two classes of interest. The data matrix ***A*** is separated into two matrices ***A***_0_ and ***A***_1_, each containing the subset of rows (representing cells) for the corresponding class of interest. For example, the rows of ***A*** representing cells in the MS lineage can be placed in ***A***_0_, and the other rows (representing cells in other lineages) can be placed in ***A***_1_. The maximum class separation occurs when the data are projected onto the vector

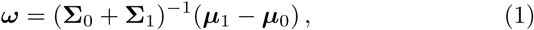

i.e. the inverse of the sum of the covariance matrices of the data matrices for each class, ***A***_0_ and ***A***_1_, multiplied by the vector difference in class means. The data matrices can be projected onto ω by simple matrix-vector multiplication (***A***_0_ ω and ***A***_1_ ω, or projected collectively as ***A ω***) to obtain a one-dimensional representation of the data points, maximally separated into the two classes.

In our use case, the matrices ***A***_0_ and ***A***_1_ have cells of their respective class in the rows and genes in the columns, i.e. they are of dimensions 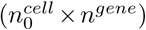 and 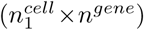 respectively. The column covariance matrices Σ_0_ and Σ_1_ are both of dimension (*n^gene^ × n^gene^*). The mean gene expression vectors µ_0_ and µ_1_ are both of dimension (*n^gene^* × 1), derived from the column means of ***A***_0_ and ***A***_1_ respectively (i.e. these vectors are the mean expression level for each gene within the class). The resulting FDA projection vector ω is of dimension (*n^gene^* × 1).

The vector ω gives the set of gene weightings that maximally separates two classes of cells when expressed as a linear combination (***A***_0_ ω and ***A***_1_ ω). Each component of ω is the weight of a specific gene, and therefore the absolute magnitude of these weights is directly related to how differentially expressed a gene is between the two classes, relative to other genes.

FDA doesn’t yield an innately optimal class separation boundary along its projection vector, ω. For Figure 4 and the figures in Table S4, we plotted as the decision boundary the line which equalized the percentage of cells in Class 1 above the line and the percentage of cells in class 0 below the line.

FDA is a particularly useful technique for identifying genes differentially involved in different developmental processes (or in other A/B testing scenarios, such as case vs. control). However, it appears that this type of discriminant analysis is not yet widely used for this purpose in the biological sciences. We provided a number of examples of how to apply this technique to a whole-organism dataset, and enhancements of this technique useful for the biological sciences, such as aggregation of GO term weights to understand the developmental roles of positively and negatively weighted genes, as observed in the resultant FDA weightings.

## Supplementary Tables

**Table S1:** The binarized gene expression matrix, with cells in the rows, genes in the columns, and 0 or 1 in the cells indicating whether or not gene expression crossed the minimum expression threshold during the lifetime of the cell.

**Table S2:** (a) The eigenvalues of the binarized gene expression matrix, with eigen-decomposition performed on the columns (i.e. the genes). (b) The corresponding eigenvectors for each eigenvalue, i.e. the principal axes of gene expression.

**Table S3:** The projection of the binarized gene expression matrix onto the principal component axes. Rows represent cells; columns represent principal component axes. The PC1, PC2 and PC3 columns are the source of the data plotted in Figure 1.

**Table S4:** The Fisher’s Discriminant Analysis (FDA) results for the projection of the gene expression profiles onto the FDA ω-vector, which is the axis that maximizes inter-class variance and minimizes intra-class variance between two classes. Numerous binary class separations are present in this result set, including cell lineages (AB, MS, C, D, E) with filenames names containing e.g. “cell-group:E_lineage”; anatomy ontology terms, including tissue types, with filenames containing “anat:”; developmental stages, with filenames containing “life-stage:”; cell division direction, with filenames containing e.g. “direction:a” for anterior or “direction:p” for posterior; cell division depth, with filenames containing “lineage-depth:”; whether a cell undergoes apoptosis, becomes part of a syncitium, or is terminally differentiated, with filenames containing “death-in:”, “syn-in:” or “is-leaf:”, followed by the gender (herm, male, both, one-or-both). Each result filename is preceded with a number, ranging from 01.09794 to 57.80538, which is the ratio of inter-class to intra-class variance, such that larger numbers indicate a cleaner separation by FDA into two compact and widely-separated classes. Each result includes three files with the same filename prefix: one file ending “.pdf”, which is a diagram showing the class separation achieved by FDA vs. the developmental birthtime of each cell; one file ending in “-gene-weights.csv”, indicating the weight of each gene, obtained by FDA analysis, where a larger negative weight indicates the gene was only active in cells that did not have the trait listed in the filename, and where a larger positive weight indicates that the gene was only active in cells that exhibited the trait listed in the filename; and one file ending “-goterm-weights.tsv” indicating the summed (total) weight of each Gene Ontology term across all cells, giving a measure of how much each gene annotated with the given GO term contributed towards the cells falling in Class 0 or Class 1.

**Table S5:** The linear weighting x of genes that maps the gene expression profile of a cell onto the developmental age of the cell with the minimum squared error. This is derived by solving the linear system ***Ax*** = ***b***, where ***A*** is the binarized gene expression matrix and ***b*** is the vector of cell birth times. Large negative and positive weights here correspond with genes that require a stronger contribution to linearize the mapping from gene expression to cell birth time.

**Table S6:** Comparison for each cell of b (cell birth time), PC1 (the projection of gene expression for the cell onto the first principal component axis), Ax (the linear combination of gene expression that best maps gene the expression profile of the cell onto the cell birth time), the Sulston cell onset time (obtained from [11], not available for all cells), the average offset from the cell birth time to the first point at which a gene’s expression levels cross the binarization threshold, the average TTL (time to live) for the cell (i.e. the time until cell division or apoptosis), and the θ- and y- coordinates for the cylindrical projection of the gene expression manifold.

**Table S7:** The projection of gene expression values onto the principal component axes for the PCA of the transpose of the binarized gene expression matrix (i.e. the PCA of the transpose of Table S1). Consequently, this represents dimension reduction in the cell dimensions rather than the gene dimensions. Whereas the first k columns of Table S3 represents a dimension-reduced gene expression profiles for each cell, the first k columns of Table S7 represents a dimension-reduced profile of gene expression patterns for each gene. Genes that have similar dimension-reduced profiles (i.e. genes that are close together in the PCA space) are expressed in many of the same cells.

## Declarations

**Ethics approval and consent to participate**

Not applicable.

## Consent for publication

Not applicable.

## Availability of data and material

All data generated during this study are included in this published article and its supplementary information files. All data analyzed during this study are publicly available in the Bibliography.

## Authors’ contributions

LH performed the data import, integration, analysis, and visualization work, and drafted the manuscript. BB and IK contributed literature search, biological context analysis, research direction, and manuscript editing.

## Acknowledgments

We thank Sumaiya Nazeen and Hoon Cho for helpful comments and suggestions.

---

**Figure S1.**
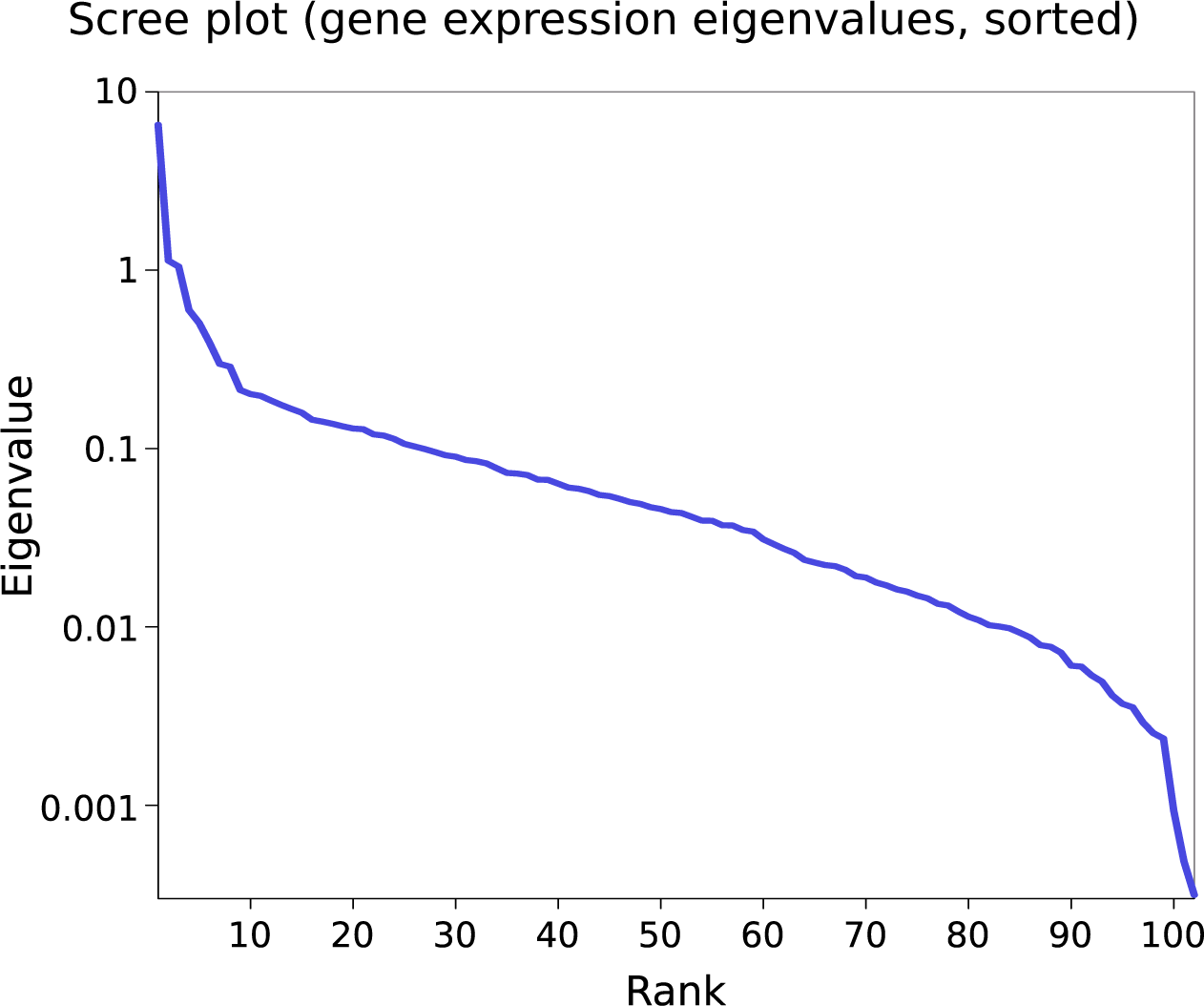
The scree plot from Principal Components Analysis of gene expression (i.e. the eigenvalues of gene expression space, sorted in decreasing order of magnitude). Note the vertical log scale. This plot shows that a significant amount of variance in gene expression across cells is captured by the first 10 components of the PCA.

**Figure S2.**
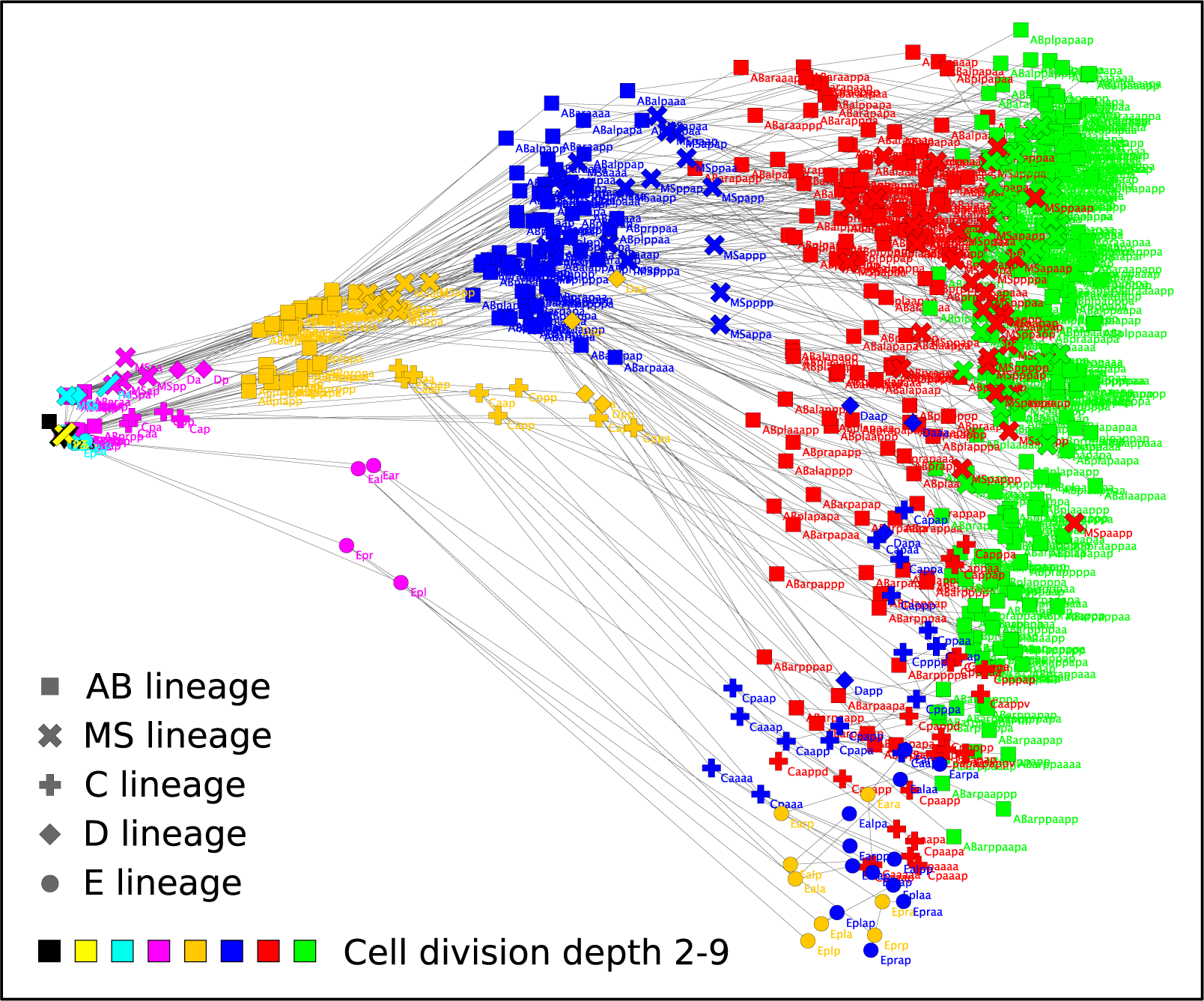
The cylindrical projection of the gene expression manifold seen in Figure 1 onto a flat 2D surface. The horizontal axis is θ, the angle of rotation about the cylindrical axis, and the vertical axis, y, represents the distance of the cell along the cylindrical axis. (The cell identities are labeled for illustration purposes only, and are not in most cases legible, due to the label size and high degree of overlap.)

**Figure S3.**
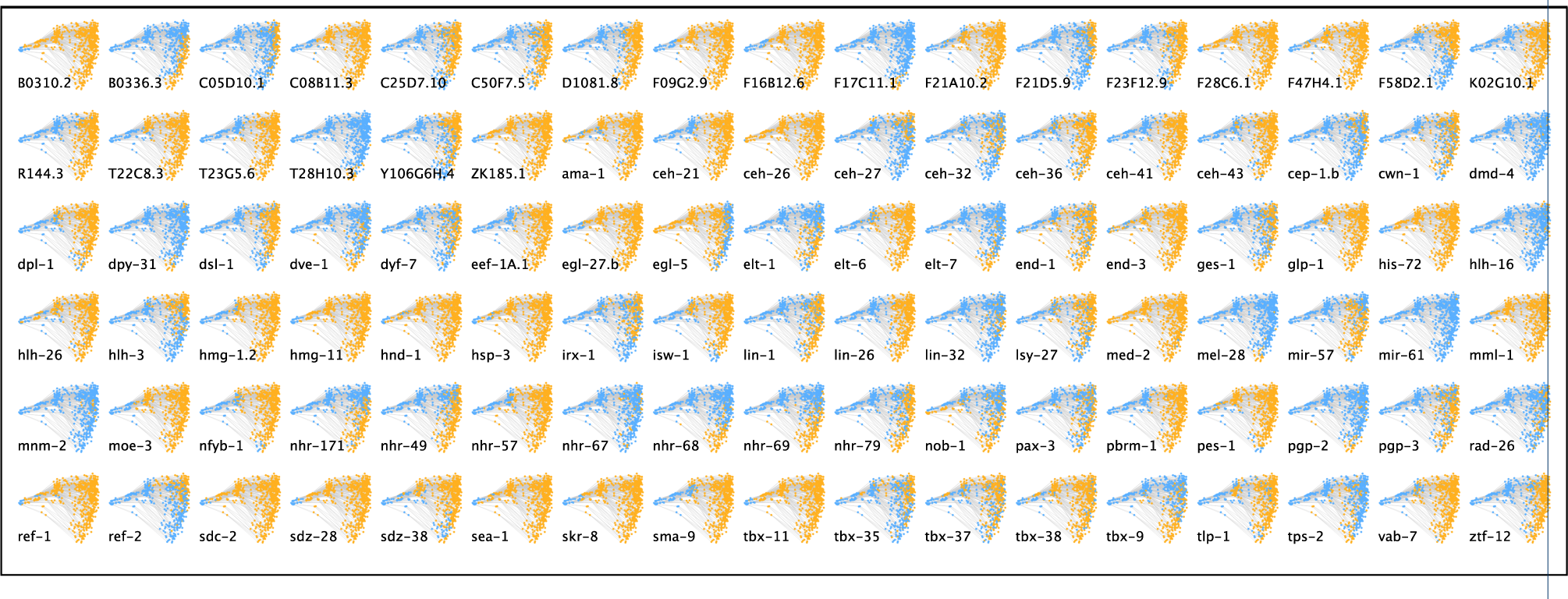
The cylindrical projection of the gene expression manifold for each of the 102 genes, with yellow and blue indicating that the gene was respectively expressed or not expressed in a given cell above the chosen expression threshold.

**Figure S4.**
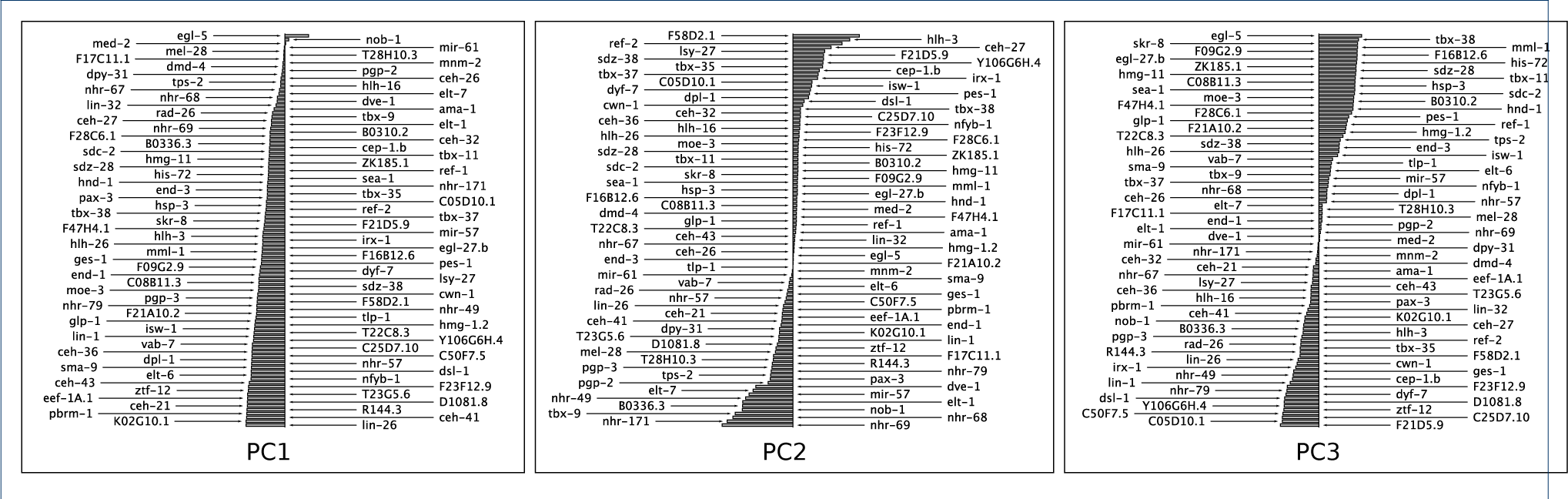
Bar chart of eigenvector components for PC1-PC3, with the components sorted vertically in order of magnitude (positive to the right of the midline, negative to the left), showing the relative magnitude of contribution of each gene to the position of cells in each of the three PC axes. Multiplying these weights by the square root of the corresponding eigenvalue gives the principle component loadings. Larger positive or negative weights indicate more relative contribution of the gene to the position of a cell in the corresponding PC axis. For PC2, only a minority of genes have large negative or positive weight, indicating this subset of genes is involved in differentiation. For PC1, the comparative lack of low-weighted genes indicates that the strong correlation between PC1 and the developmental age of the organism is not a simple relationship governed by a small subset of genes, but that most of the genes are involved in this correlation.

## References

1. Moore, J.L., Du, Z., Bao, Z.: Systematic quantification of developmental phenotypes at single-cell resolution during embryogenesis. Development 140(15), 3266–3274 (2013)

2. Richards, J.L., Zacharias, A.L., Walton, T., Burdick, J.T., Murray, J.I.: A quantitative model of normal *Caenorhabditis elegans* embryogenesis and its disruption after stress. Developmental biology 374(1), 12–23 (2013)

3. Giurumescu, C.A., Kang, S., Planchon, T.A., Betzig, E., Bloomekatz, J., Yelon, D., Cosman, P., Chisholm, A.D.: Quantitative semi-automated analysis of morphogenesis with single-cell resolution in complex embryos. Development 139(22), 4271–4279 (2012)

4. Liu, X., Long, F., Peng, H., Aerni, S.J., Jiang, M., Sánchez-Blanco, A., Murray, J.I., Preston, E., Mericle, B., Batzoglou, S., et al.: Analysis of cell fate from single-cell gene expression profiles in *C. elegans*. Cell 139(3), 623–633 (2009)

5. Bao, Z., Murray, J.I., Boyle, T., Ooi, S.L., Sandel, M.J., Waterston, R.H.: Automated cell lineage tracing in *Caenorhabditis elegans*. Proceedings of the National Academy of Sciences of the United States of America 103(8), 2707–2712 (2006)

6. Boyle, T.J., Bao, Z., Murray, J.I., Araya, C.L., Waterston, R.H.: AceTree: a tool for visual analysis of *Caenorhabditis elegans* embryogenesis. BMC bioinformatics 7(1), 275 (2006)

7. Murray, J.I., Bao, Z., Boyle, T.J., Waterston, R.H.: The lineaging of fluorescently-labeled *Caenorhabditis elegans* embryos with StarryNite and AceTree. Nature protocols 1(3), 1468–1476 (2006)

8. Murray, J.I., Bao, Z., Boyle, T.J., Boeck, M.E., Mericle, B.L., Nicholas, T.J., Zhao, Z., Sandel, M.J., Waterston, R.H.: Automated analysis of embryonic gene expression with cellular resolution in *C. elegans*. Nature methods 5(8), 703–709 (2008)

9. Cao, J., Packer, J.S., Ramani, V., Cusanovich, D.A., Huynh, C., Daza, R., Qiu, X., Lee, C., Furlan, S.N., Steemers, F.J., Adey, A., Waterston, R.H., Trapnell, C., Shendure, J.: Comprehensive single cell transcriptional profiling of a multicellular organism by combinatorial indexing. Science 357, 661–667 (2017)

10. Cho, H., Berger, B., Peng, J.: Neural data visualization for scalable and generalizable single cell analysis. Cell Systems (2018)

11. Murray, J.I., Boyle, T.J., Preston, E., Vafeados, D., Mericle, B., Weisdepp, P., Zhao, Z., Bao, Z., Boeck, M., Waterston, R.H.: Multidimensional regulation of gene expression in the *C. elegans* embryo. Genome Research 22(7), 1282–1294 (2012)

12. Waterston et al.: Expression Patterns in Caenorhabditis (EPIC). http://epic.gs.washington.edu/ Accessed 2018-06-28

13. Cooke, J.: Control of somite number during morphogenesis of a vertebrate, Xenopus laevis (1975)

14. Dale, K.J., Pourquie, O.: A clock-work somite. Bioessays 22(1), 72–83 (2000)

15. Lorthongpanich, C., Doris, T.P.Y., Limviphuvadh, V., Knowles, B.B., Solter, D.: Developmental fate and lineage commitment of singled mouse blastomeres. Development 139(20), 3722–3731 (2012)

16. Desai, A.R., McConnell, S.K.: Progressive restriction in fate potential by neural progenitors during cerebral cortical development. Development 127(13), 2863–2872 (2000)

17. Palmeirim, I., Henrique, D., Ish-Horowicz, D., Pourquié, O.: Avian *hairy* gene expression identifies a molecular clock linked to vertebrate segmentation and somitogenesis. Cell 91(5), 639–648 (1997)

18. Satoh, N.: Timing mechanisms in early embryonic development. Differentiation 22(1-3), 156–163 (1982)

19. Reinhart, B.J., Slack, F.J., Basson, M., Pasquinelli, A.E., Bettinger, J.C., Rougvie, A.E., Horvitz, H.R., Ruvkun, G.: The 21-nucleotide *let-7* RNA regulates developmental timing in *Caenorhabditis elegans*. nature 403(6772), 901–906 (2000)

20. Zhao, Z., Boyle, T.J., Liu, Z., Murray, J.I., Wood, W.B., Waterston, R.H.: A negative regulatory loop between microRNA and *Hox* gene controls posterior identities in *Caenorhabditis elegans*. PLoS Genet 6(9), 1001089 (2010)

21. Hench, J., Henriksson, J., Abou-Zied, A.M., Lüppert, M., Dethlefsen, J., Mukherjee, K., Tong, Y.G., Tang, L., Gangishetti, U., Baillie, D.L., et al.: The homeobox genes of *Caenorhabditis elegans* and insights into their spatio-temporal expression dynamics during embryogenesis. PloS one 10(5), 0126947 (2015)

22. Nair, G., Walton, T., Murray, J.I., Raj, A.: Gene transcription is coordinated with, but not dependent on, cell divisions during *C. elegans* embryonic fate specification. Development 140(16), 3385–3394 (2013)

23. Treutlein, B., Brownfield, D.G., Wu, A.R., Neff, N.F., Mantalas, G.L., Espinoza, F.H., Desai, T.J., Krasnow, M.A., Quake, S.R.: Reconstructing lineage hierarchies of the distal lung epithelium using single-cell RNA-seq. Nature 509(7500), 371–375 (2014)

24. Yan, L., Yang, M., Guo, H., Yang, L., Wu, J., Li, R., Liu, P., Lian, Y., Zheng, X., Yan, J., et al.: Single-cell RNA-Seq profiling of human preimplantation embryos and embryonic stem cells. Nature structural & molecular biology 20(9), 1131–1139 (2013)

25. Trapnell, C., Cacchiarelli, D., Grimsby, J., Pokharel, P., Li, S., Morse, M., Lennon, N.J., Livak, K.J., Mikkelsen, T.S., Rinn, J.L.: The dynamics and regulators of cell fate decisions are revealed by pseudotemporal ordering of single cells. Nature biotechnology 32(4), 381–386 (2014)

26. Bendall, S.C., Davis, K.L., Amir, E.-a.D., Tadmor, M.D., Simonds, E.F., Chen, T.J., Shenfeld, D.K., Nolan, G.P., Pe’er, D.: Single-cell trajectory detection uncovers progression and regulatory coordination in human B cell development. Cell 157(3), 714–725 (2014)

27. Davis, K.L., Bendall, S.C., El-ad, D.A., Simonds, E.F., Jager, A., Trejo, A., Pe’er, D., Nolan, G.P.: Single cell trajectory detection orders hallmarks of early human B cell development. Blood 120(21), 1044–1044 (2012)

28. Araya, C.L., Kawli, T., Kundaje, A., Jiang, L., Wu, B., Vafeados, D., Terrell, R., Weissdepp, P., Gevirtzman, L., Mace, D., et al.: Regulatory analysis of the *C. elegans* genome with spatiotemporal resolution. Nature 512(7515), 400–405 (2014)

29. Wagner, D.E., Weinreb, C., Collins, Z.M., Briggs, J.A., Megason, S.G., Klein, A.M.: Single-cell mapping of gene expression landscapes and lineage in the zebrafish embryo. Science (2018)

30. Farrell, J.A., Wang, Y., Riesenfeld, S.J., Shekhar, K., Regev, A., Schier, A.F.: Single-cell reconstruction of developmental trajectories during zebrafish embryogenesis. Science (2018)

31. Briggs, J.A., Weinreb, C., Wagner, D.E., Megason, S., Peshkin, L., Kirschner, M.W., Klein, A.M.: The dynamics of gene expression in vertebrate embryogenesis at single-cell resolution. Science (2018)

32. Lee, M.T., Bonneau, A.R., Giraldez, A.J.: Zygotic genome activation during the maternal-to-zygotic transition. Annual review of cell and developmental biology 30, 581–613 (2014)

33. Goszczynski, B., McGhee, J.D.: Reevaluation of the role of the *med-1* and *med-2* genes in specifying the *Caenorhabditis elegans* endoderm. Genetics 171(2), 545–555 (2005)

34. Nicholas, H.R., Hodgkin, J.: The *C. elegans Hox* gene *egl-5* is required for correct development of the hermaphrodite hindgut and for the response to rectal infection by Microbacterium nematophilum. Developmental biology 329(1), 16–24 (2009)

35. Robertson, S.M., Medina, J., Lin, R.: Uncoupling Different Characteristics of the *C. elegans* E Lineage from Differentiation of Intestinal Markers. PLoS ONE 9 (2014)

36. Zhu, J., Fukushige, T., McGhee, J.D., Rothman, J.H.: Reprogramming of early embryonic blastomeres into endodermal progenitors by a *Caenorhabditis elegans GATA* factor. Genes Dev. 12, 3809–3814 (1998)

37. Van Auken, K., Weaver, D.C., Edgar, L.G., Wood, W.B.: *Caenorhabditis elegans* embryonic axial patterning requires two recently discovered posterior-group *Hox* genes. Proceedings of the National Academy of Sciences 97(9), 4499–4503 (2000)

38. Shao, J., He, K., Wang, H., Ho, W.S., Ren, X., An, X., Wong, M.K., Yan, B., Xie, D., Stamatoyannopoulos, J., et al.: Collaborative regulation of development but independent control of metabolism by two epidermis-specific transcription factors in *Caenorhabditis elegans*. Journal of Biological Chemistry 288(46), 33411–33426 (2013)

39. Chen, Z., Eastburn, D.J., Han, M.: The *Caenorhabditis elegans* nuclear receptor gene *nhr-25* regulates epidermal cell development. Molecular and cellular biology 24(17), 7345–7358 (2004)

40. Thompson, K.W., Joshi, P., Dymond, J.S., Gorrepati, L., Smith, H.E., Krause, M.W., Eisenmann, D.M.: The paired-box protein PAX-3 regulates the choice between lateral and ventral epidermal cell fates in *C. elegans*. Developmental biology 412(2), 191–207 (2016)

41. Andachi, Y.: *Caenorhabditis elegans* T-box genes *tbx-9* and *tbx-8* are required for formation of hypodermis and body-wall muscle in embryogenesis. Genes to Cells 9(4), 331–344 (2004)

42. Anavy, L., Levin, M., Khair, S., Nakanishi, N., Fernandez-Valverde, S.L., Degnan, B.M., Yanai, I.: BLIND ordering of large-scale transcriptomic developmental timecourses. Development 141(5), 1161–1166 (2014)

43. Hendriks, G.-J., Gaidatzis, D., Aeschimann, F., Großhans, H.: Extensive oscillatory gene expression during *C. elegans* larval development. Molecular cell 53(3), 380–392 (2014)

44. Kim, D.h., Grün, D., van Oudenaarden, A.: Dampening of expression oscillations by synchronous regulation of a microRNA and its target. Nature genetics 45(11), 1337–1344 (2013)

45. McCulloch, K.A., Rougvie, A.E.: *Caenorhabditis elegans* period homolog *lin-42* regulates the timing of heterochronic miRNA expression. Proceedings of the National Academy of Sciences 111(43), 15450–15455 (2014)

46. Gorban, A.N., Zinovyev, A.Y.: Principal graphs and manifolds. In: Olivas, E.S. (ed.) Handbook of Research on Machine Learning Applications and Trends: Algorithms, Methods, and Techniques, pp. 28–59. IGI Global, Pennsylvania, USA (2009)

